# Pathogenesis-related protein 1 (PR-1) genes in soybean: genome-wide identification, structural analysis and expression profiling under multiple biotic and abiotic stresses

**DOI:** 10.1101/2021.03.27.437342

**Authors:** Fabricio Almeida-Silva, Thiago M. Venancio

**Affiliations:** Laboratório de Química e Função de Proteínas e Peptídeos, Centro de Biociências e Biotecnologia, Universidade Estadual do Norte Fluminense Darcy Ribeiro, Campos dos Goytacazes, RJ, Brazil

**Keywords:** plant immunity, phytopathology, gene family evolution, functional genomics

## Abstract

Plant pathogenesis-related (PR) proteins are a large group of proteins, classified in 17 families, that are induced by pathological conditions. Here, we characterized the soybean PR-1 (GmPR-1) gene repertoire at the sequence, structural and expression levels. We found 24 GmPR-1 genes, clustered in two phylogenetic groups. GmPR-1 genes are under strong purifying selection, particularly those that emerged by tandem duplications. GmPR-1 promoter regions are abundant in cis-regulatory elements associated with major stress-related transcription factor families, namely WRKY, ERF, HD-Zip, C2H2, NAC, and GATA. We observed that 23 GmPR-1 genes are induced by stress conditions or exclusively expressed upon stress. We explored 1972 transcriptome samples, including 26 stress conditions, revealing that most GmPR-1 genes are differentially expressed in a plethora of biotic and abiotic stresses. Our findings highlight stress-responsive GmPR-1 genes with potential biotechnological applications, such as the development of transgenic lines with increased resistance to biotic and abiotic stresses.

## 1 Introduction

Plants have evolved a complex genetic machinery to cope with pathogens and pests. Plant pathogenesis-related (PR) proteins are core components of plant defense and are classified in 17 families based on their sequence similarities, enzymatic activities and structural properties [1]. PR-1, the first identified family of PR proteins, was discovered in tobacco leaves infected with tobacco mosaic virus [2]. Alexander *et al.* [2] reported that accumulated PR-1 proteins comprised 2% of the total proteins in infected tobacco leaves, suggesting a prominent role in plant defense.

PR-1 proteins are ubiquitous and have been identified in many plant species [3–6]. They can be acidic or alkaline and are mostly secreted to the apoplast, although some can be stored in vacuoles [7]. All PR-1 proteins share a conserved CAP (cysteine-rich secretory protein, antigen 5, pathogenesis-related 1) domain that typically folds into 4 α-helices and 4-βsheets stabilized by disulfide bonds [8]. These unique features are responsible for their biological functions, which are mainly associated with defense to biotic and abiotic stresses [4,9]. However, some works have also linked PR-1 proteins with endogenous functions, such as leaf senescence and floral development [10].

Soybean (*Glycine max* (L.) Merr.) is the most important legume crop worldwide, being used in human and animal nutrition, and industrial applications. Soybean diseases are responsible for an economic loss of 4.6 billion dollars only in the US [11]. Likewise, soybean yield is significantly affected by multiple environmental factors [12]. Transgenic plants overexpressing PR-1 genes have demonstrated increased resistance to both biotic and abiotic stresses [5,13,14]. However, the soybean PR-1 gene repertoire has never been systematically characterized. Hence, a comprehensive analysis of the soybean PR-1 genes and their transcriptional profiles under multiple stresses would certainly be useful to pinpoint potential targets for biotechnological applications, such as developing transgenic lines with increased resistance.

Here, we have characterized the soybean PR-1 (GmPR-1) gene repertoire at the sequence, structural and transcriptional levels, under diverse tissues and stress conditions. We found 24 GmPR-1 genes clustered in two phylogenetic groups. All GmPR-1 genes are under strong purifying selection, particularly tandem-derived paralogs, and they tend to have tissue-specific expression. Besides, we observed that major stress-related transcription factor (TF) families (*e.g.,* WRKY, C2H2, and ERF) are putative regulators of GmPR-1 gene expression. Finally, we demonstrated that most GmPR-1 genes are transcriptionally modulated by a plethora of biotic and abiotic stresses, from which one is exclusively expressed upon stress. We reported the transcriptional landscape of GmPR-1 genes under 26 different biotic and abiotic stress conditions, making this study the most comprehensive investigation of GmPR-1 transcriptional response to stresses. Our findings can help select potential GmPR-1 genes to be used in the development of transgenic lines with increased stress resistance.

## 2 Materials and Methods

### 2.1 Identification of GmPR-1 genes

Soybean predicted proteins were downloaded from the PLAZA database [15]. GmPR-1 proteins were identified with BLASTp searches [16] against the soybean predicted proteins (E-value threshold: 1e-10) using curated *Arabidopsis thaliana* PR-1 proteins as queries (UniProt IDs: P33154 and Q9ZNS4). The proteins found in the BLASTp search were then scanned for the presence of the CAP domain (PF00188). Further, we confirmed that all GmPR-XS1 genes were included in the same subfamily in PLAZA (ORTHO04D000086) [15]. Functional gene annotations (Gene Ontology and PFAM) were retrieved from SoyBase [17] and PLAZA [15].

### 2.2 Homology modelling and structural analysis

Tertiary protein structures and the percentage of secondary structures were predicted with the Phyre2 server [18]. Physicochemical properties were identified with the R package Peptides [19]. Signal peptides were predicted with SignalP 5.0 [20]. The .gff file containing exon-intron boundaries of GmPR-1 genes was downloaded from PLAZA [15] and visualized with gggenes [21]. Protein length calculation and sequence manipulations were performed with Biostrings [22].

### 2.3 Phylogenetic reconstruction, gene duplication and selection analyses

Protein sequences were aligned with MAFFT [23] and visualized with msa [24]. Maximum likelihood phylogenetic trees were inferred using IQTREE2 [25] with 1000 bootstraps. A BLASTp search against the UniProt database [26] was performed to identify an outgroup for the tree of GmPR-1 genes, and a *Cajanus cajan* PR-1 (UniProt ID: A0A151S132) was selected as the best hit after excluding *G. max* hits. PR-1 proteins from *A. thaliana* and *Vigna radiata* were retrieved from PLAZA by searching for genes in the same subfamily of GmPR-1 genes (ORTHO04D000086). The R package ggtree [27] was used for tree visualization. Paralogous gene pairs and their ratios of the number of nonsynonymous substitutions per nonsynonymous substitution site to the number of synonymous substitutions per synonymous substitution site (Ka/Ks) were retrieved from a recent publication from our group [28]. Divergence times (in million years ago, mya) were calculated using DT = Ks / (2 × 6.1 × 1e−9) × 1e−6 mya [29].

### 2.4 Identification of conserved motifs and cis-regulatory elements

Conserved motifs were identified *de novo* with the MEME [30] algorithm implemented in the R package universalmotif [31] (nmotifs = 10, minw = 6, maxw = 50). Position weight matrices of known cis-regulatory elements (CREs) were downloaded from PlantPAN 3.0 [32]. Promoters were defined as the regions between −1000 bp and +200 bp relative to the transcription start site, as established for *A. thaliana* [33]. Promoter sequences were scanned for the presence of CREs using the function scan_sequences() from the R package universalmotif [31] with a log-odds threshold of 0.9, and TF families associated with each CRE were retrieved from PlantTFDB [34].

### 2.5 Gene expression analysis

Raw read counts were downloaded from the Soybean Expression Atlas (http://venanciogroup.uenf.br/cgi-bin/gmax_atlas/index.cgi) [35]. Samples from callus, seedling, endosperm, inflorescences and suspensor were not included in the global expression analysis because they were represented by too few samples (*n* <10). Additional 98 publicly available RNA-seq BioProjects investigating soybean response to biotic or abiotic stresses (930 samples) were downloaded from NCBI’s Sequence Read Archive using our pipeline described in [35] (Supplementary Table S1). A total of 145 pairwise comparisons for differential gene expression were performed with DESeq2 [36] (Supplementary Table S2). Genes were considered differentially expressed if the absolute fold change was greater than 1.5 and Benjamini-Hochberg-adjusted *P*-value lower than 0.05. The pairwise comparisons only included samples from the same BioProject to avoid potential artifacts that could bias biological interpretation, such as different time points, cultivars or plant age.

## 3 Results and discussion

### 3.1 Identification, distribution and physicochemical properties of GmPR-1 genes and their protein products

Using *A. thaliana* PR-1 proteins as queries, we identified 24 soybean genes encoding PR-1 proteins through BLASTp searches, all with a conserved CAP domain (PF00188). Genes were named using the standard nomenclature for other species (GmPR-1-1 to GmPR-1-24; Table 1). Most of the GmPR-1 genes were located on chromosomes 13 and 15 (*n* = 9 and 6, respectively), while others were located on chromosomes 7, 17, 1, 2, 10, and 16 (*n* = 3, 2, 1, 1, 1, 1, respectively). PR-1 genes have been found in widely variable numbers across species, such as tomato [37], grape [38], wheat [39] and rice [40], suggesting that a number of lineage-specific expansions and contractions shaped the PR-1 repertoires across species.

**Table 1.** Structural properties of the identified GmPR-1 genes.

| Gene ID | Gene name | Exon<br>no. | Length<br>(aa) | pI | MW<br>(kDa) | H | SL | SP | AH<br>% | BS<br>% | D<br>% | TH<br>% |
| --- | --- | --- | --- | --- | --- | --- | --- | --- | --- | --- | --- | --- |
| Glyma.01G180400 | GmPR-1-1 | 3 | 275 | 7.2 | 30 | -0.17 | Apoplast | - | 26 | 21 | 28 | 0 |
| Glyma.02G060800 | GmPR-1-2 | 1 | 172 | 7.4 | 19 | -0.21 | Apoplast | 21-22 | 35 | 16 | 22 | 0 |
| Glyma.07G186200 | GmPR-1-3 | 1 | 165 | 6 | 18 | -0.38 | Apoplast | 25-26 | 38 | 19 | 19 | 10 |
| Glyma.07G186300 | GmPR-1-4 | 1 | 160 | 4.9 | 18 | -0.19 | Apoplast | 23-24 | 39 | 18 | 19 | 0 |
| Glyma.07G186400 | GmPR-1-5 | 2 | 170 | 9.1 | 19 | -0.41 | Apoplast | - | 36 | 13 | 28 | 9 |
| Glyma.10G047000 | GmPR-1-6 | 2 | 277 | 8.5 | 32 | -0.84 | Apoplast | 22-23 | 22 | 13 | 48 | 0 |
| Glyma.13G094100 | GmPR-1-7 | 1 | 200 | 6.9 | 23 | -0.36 | Apoplast | 22-23 | 32 | 16 | 32 | 8 |
| Glyma.13G094200 | GmPR-1-8 | 1 | 175 | 9.1 | 20 | -0.37 | Apoplast | 25-26 | 39 | 15 | 25 | 9 |
| Glyma.13G251600 | GmPR-1-9 | 1 | 174 | 4.3 | 19 | -0.15 | Apoplast | 27-28 | 38 | 17 | 21 | 9 |
| Glyma.13G251700 | GmPR-1-10 | 1 | 174 | 4.3 | 19 | -0.15 | Apoplast | 27-28 | 38 | 17 | 21 | 9 |
| Glyma.13G252000 | GmPR-1-11 | 1 | 174 | 4.3 | 19 | -0.15 | Apoplast | 27-28 | 38 | 17 | 21 | 9 |
| Glyma.13G252300 | GmPR-1-12 | 1 | 174 | 4.3 | 19 | -0.15 | Apoplast | 27-28 | 38 | 17 | 21 | 9 |
| Glyma.13G252400 | GmPR-1-13 | 1 | 174 | 4.3 | 19 | -0.15 | Apoplast | 27-28 | 38 | 17 | 21 | 9 |
| Glyma.13G252500 | GmPR-1-14 | 1 | 79 | 5.5 | 8 | -0.05 | Apoplast | 25-26 | 59 | 3 | 23 | 0 |
| Glyma.13G252600 | GmPR-1-15 | 1 | 161 | 6.1 | 18 | -0.32 | Apoplast | 25-26 | 40 | 16 | 14 | 10 |
| Glyma.15G062300 | GmPR-1-16 | 1 | 161 | 5.8 | 18 | -0.34 | Apoplast | 25-26 | 40 | 17 | 17 | 10 |
| Glyma.15G062400 | GmPR-1-17 | 1 | 164 | 5.1 | 18 | -0.18 | Apoplast | 26-27 | 39 | 21 | 15 | 10 |
| Glyma.15G062500 | GmPR-1-18 | 1 | 164 | 8.4 | 18 | -0.26 | Apoplast | 26-27 | 38 | 20 | 15 | 10 |
| Glyma.15G062700 | GmPR-1-19 | 1 | 165 | 8 | 18 | -0.33 | Apoplast | 27-28 | 39 | 18 | 17 | 10 |
| Glyma.15G062800 | GmPR-1-20 | 1 | 174 | 4.4 | 19 | -0.06 | Apoplast | 27-28 | 40 | 17 | 20 | 9 |
| Glyma.15G062900 | GmPR-1-21 | 1 | 126 | 4.4 | 14 | -0.4 | Apoplast | - | 35 | 25 | 12 | 0 |
| Glyma.16G143300 | GmPR-1-22 | 1 | 174 | 8.8 | 19 | -0.11 | Apoplast | 21-22 | 36 | 16 | 21 | 9 |
| Glyma.17G066000 | GmPR-1-23 | 1 | 168 | 8.7 | 19 | -0.45 | Apoplast | 17-18 | 37 | 17 | 22 | 0 |
| Glyma.17G066100 | GmPR-1-24 | 1 | 123 | 6.9 | 15 | -0.61 | Apoplast | - | 43 | 11 | 19 | 0 |

H: GRAVY hydrophobicity index based on the Kyte-Doolittle scale; SL: Subcellular localization; SP: Signal peptide cleavage site; AH, BS, D and TH: Predicted percentage of alfa-helices, beta-sheets, disordered regions and transmembrane helices in the protein structure.

The stress-related functions of GmPR-1 genes are also supported by their associated Gene Ontology terms, which are all related to defense against biotic and abiotic stresses (Supplementary Table S3). A structural analysis revealed that GmPR-1 genes have 8-32 kDa, 79-277 amino acid residues, and their isoelectric points range from 4.3 to 9.1, with 9 GmPR-1 proteins being alkaline and 15 acidic. All GmPR-1 proteins are hydrophilic (GRAVY index <0) and predicted to be secreted to the apoplast, although the signal peptide was not detectable in four of them (Table 1). The median percentage of alpha helices, beta sheets, disordered regions and transmembrane helices were 38, 17, 21, and 9, respectively. Overall, our findings are in line with those previously reported for other species [8,41].

### 3.2 A deletion on the CAP domain removed the caveolin-binding motif and the CAPE peptide in a GmPR-1 gene

The multiple sequence alignment of GmPR-1 proteins reveal a high conservation of the CAP domain across sequences (Fig. 1a). The tertiary structures of the GmPR-1 proteins are also conserved, in particular regarding the α-β-α sandwich that is distinctive of PR-1 proteins (Fig. 1b) [8]. However, *GmPR-1-14* lost a large region of the CAP domain, including the caveolin-binding motif (CBM, responsible for the sterol-binding activity) and the C-terminal region that can be cleaved to release the CAP-derived peptide (CAPE). Expectedly, this deletion significantly affected the tertiary structure of the GmPR-1-14 protein, which differs from the canonical α-β-α sandwich and folds into 2 α-helices linked by a loop (Fig. 1b).

**Fig. 1.**
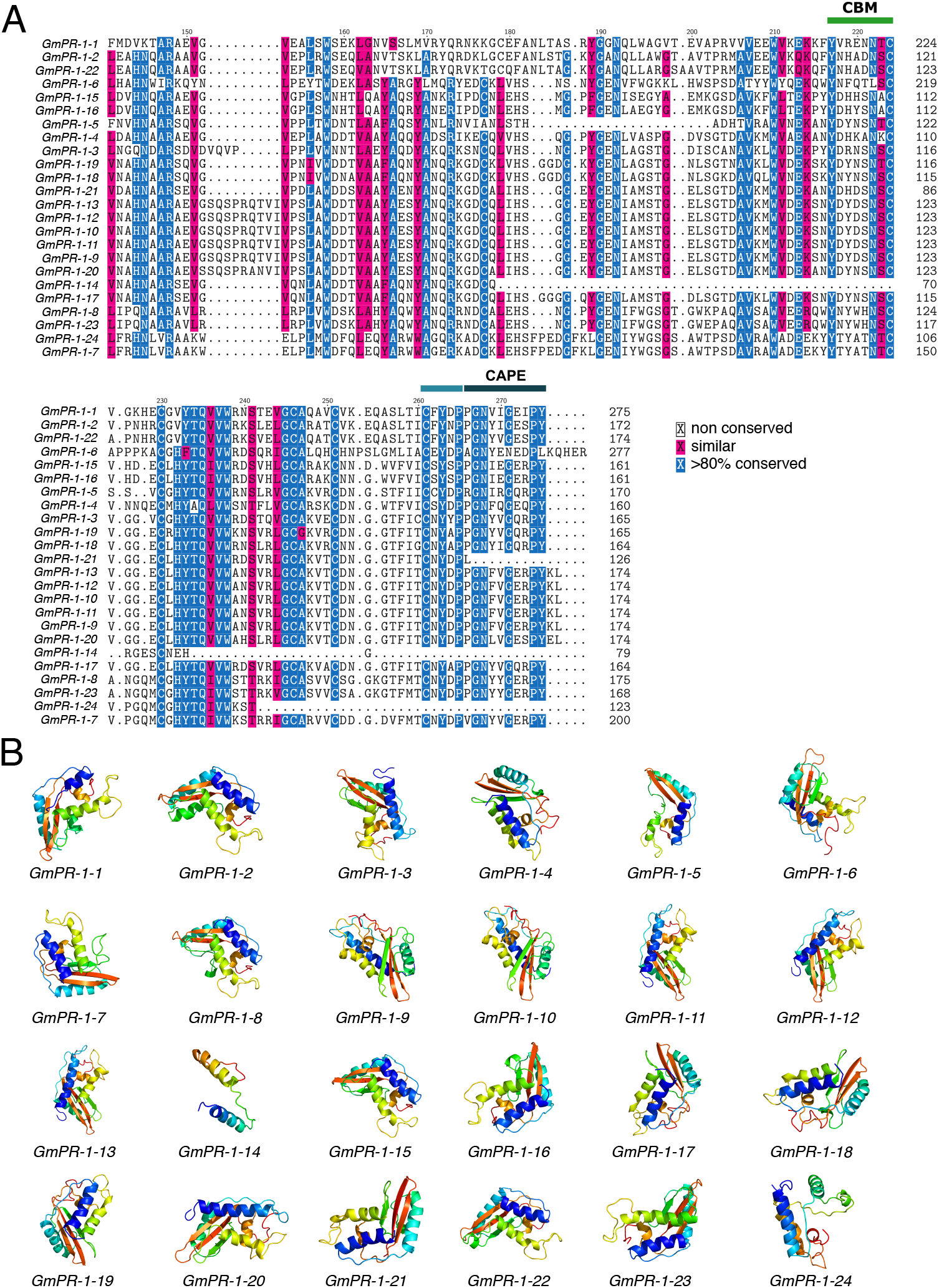
Multiple sequence alignment and predicted tertiary structures of GmPR-1 proteins. (A) Multiple sequence alignment of the CAP domain. CBM, caveolin-binding motif (responsible for sterol binding). CAPE, CAP-derived peptide. Light blue, CAPE cleavage motif. (B) Tertiary structures of GmPR-1 proteins in amino to carboxy rainbow color key. Blue represents N-terminus and red represents C-terminus. For the correspondence between gene IDs (Wm82.a2.v1) and gene names, see Table 1.

The antimicrobial activity of PR-1 proteins has been linked to their ability to bind and sequester sterols from the pathogen’s membrane [42]. This theory is supported by the observation that PR-1 proteins are especially effective against sterol auxotrophs, which are unable to produce their own sterols and depend on sterols from the environment [10]. Additionally, the last 11 amino acid residues of the CAP domain can be cleaved and release a CAP-derived peptide (CAPE), which can act indirectly in plant defense by inducing the expression of several defense-related genes [43]. As *GmPR-1-14* lacks both the CBM and the CAPE peptide, this gene has likely lost its antimicrobial properties or it has undergone functional divergence and act in plant defense by some undescribed mechanism.

### 3.3 GmPR-1 genes cluster in two phylogenetic groups

We reconstructed the phylogeny of PR-1 genes in soybean, *Vigna radiata* and *A. thaliana* (*n*=24, 8 and 22, respectively) to understand the evolutionary history of GmPR-1 genes (Fig. 2). PR-1 genes from the three species clustered into 3 groups. Cluster 1 comprised the majority of GmPR-1 genes (*n*=13), while cluster 2 and 3 contained 2 and 8 GmPR-1 genes, respectively. Cluster 1 comprises mainly genes that arose from recent local duplications, as revealed by their short branch lengths and close genomic location. A similar pattern was observed for AtPR-1 genes, with 4 genes derived from local duplications in cluster 1. Additionally, most of the GmPR-1 genes in clusters 2 and 3 arose from speciation events in the legume clade, as opposed to cluster 1, which is dominated by expansions in the soybean lineage after the split from *V. radiata*.

**Fig. 2.**
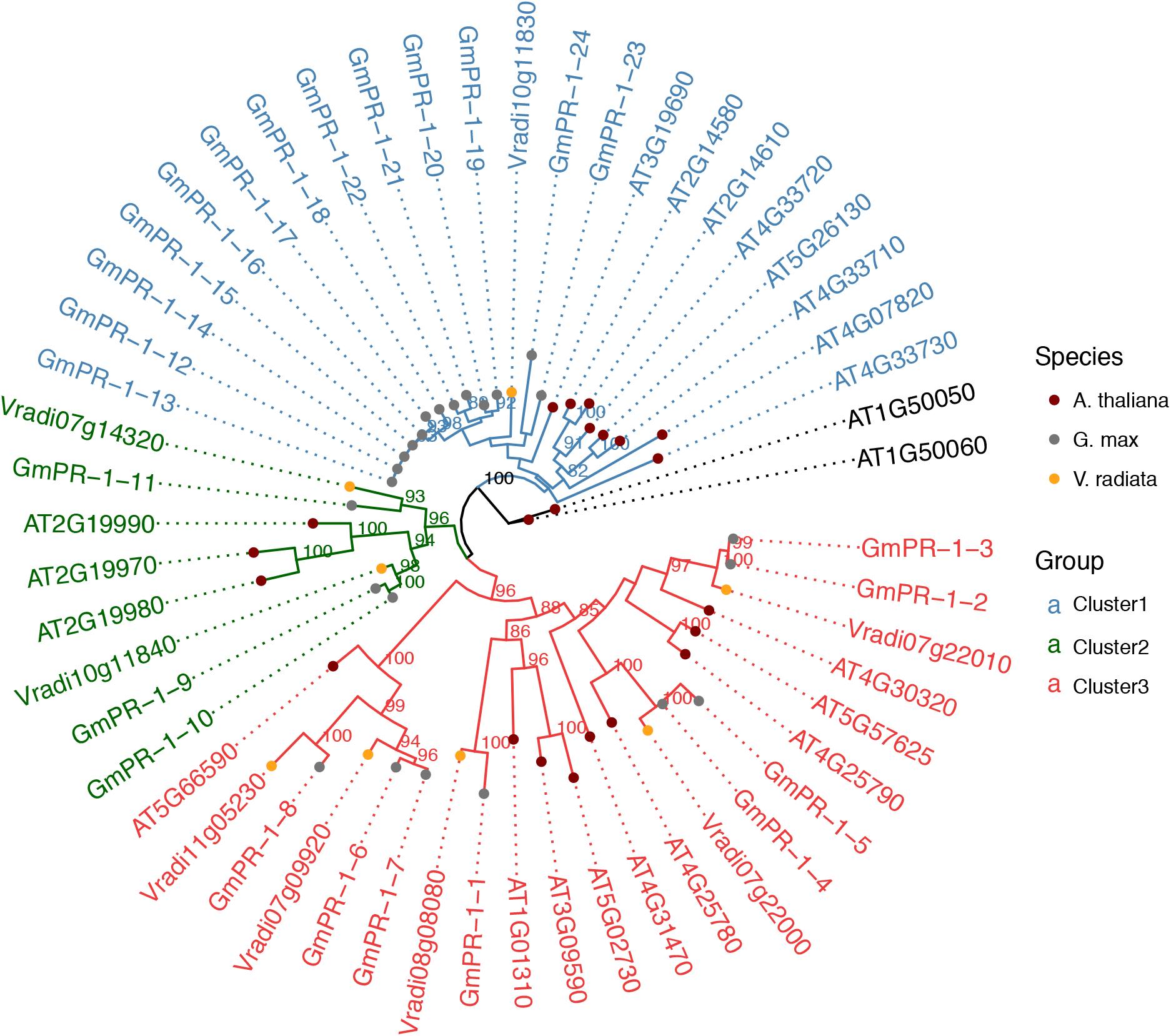
Phylogenetic reconstruction of PR-1 proteins from *Glycine max, Vigna radiata* and *Arabidopsis thaliana*. The unrooted phylogenetic tree was inferred with IQTREE2. Bootstrap support values >80 are shown in nodes. Different species and different phylogenetic clusters are represented by distinct tip colors and branch colors, respectively.

The maximum likelihood phylogenetic tree of GmPR-1 genes was reconstructed using a *C. cajan* protein (UniProt ID: A0A151S132) as outgroup. The GmPR-1 genes clustered in two groups (Fig. 3a), as observed for grapevine [38] and tomato [37], but not for wheat [39] and rice [40], which were divided in three clusters. The only three GmPR-1 genes that contain introns belong to cluster 1 (Fig. 3b), although these introns were likely acquired independently, as they are in different terminal branches of the phylogenetic tree. The same trend was observed for the position and presence/absence of motifs, from which some are exclusive to cluster 1, and others co-located near the N-terminus in cluster 2 (Fig. 3c). We also observed that GmPR-1 genes in cluster 2 mostly arose from recent local duplication events, as demonstrated by their shorter branch lengths in comparison to cluster 1.

**Fig. 3.**
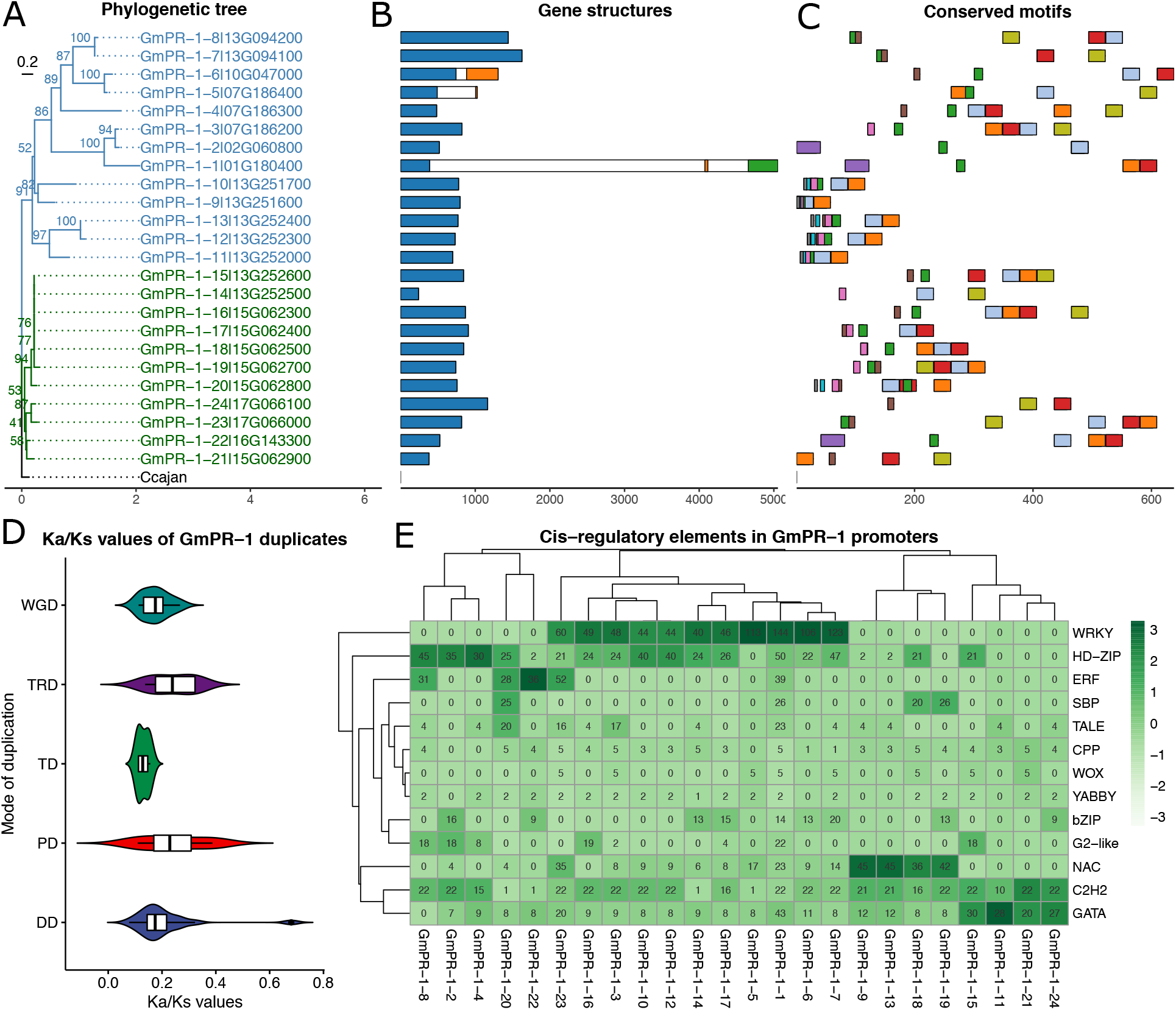
Genomic properties of GmPR-1 genes. (A) Maximum-likelihood tree inferred with IQTREE2, with 1000 bootstraps and a *Cajanus cajan* sequence (UniProt ID: A0A151S132) used as outgroup. Bootstrap support values are shown in nodes. Partial Wm82.a2.v1 gene IDs are represented after each gene’s name to emphasize the close genomic positions. (B) Gene structures demonstrating exon-intron boundaries. (C) Conserved motifs identified *de novo* with MEME. (D) Ratio of the number of nonsynonymous substitutions per nonsynonymous substitution site (Ka) to the number of synonymous substitutions per synonymous substitution site (Ks). WGD, whole-genome duplication. TRD, transposed duplication. TD, tandem duplication. PD, proximal duplication. DD, dispersed duplication. (E) Heatmap of absolute frequency of known *cis*-regulatory elements (CREs) in promoter sequences of GmPR-1 genes. Genes and samples were hierarchically clustered based on the matrix of CRE frequencies. Color intensities are scaled by column to highlight the most abundant CREs for each GmPR-1 gene.

### 3.4 Evolution of GmPR-1 genes is constrained by strong purifying selection

All GmPR-1 duplicated genes were classified into 5 modes of duplication according to the algorithm described in [44]: dispersed, proximal, tandem, transposed and whole-genome duplications (DD, PD, TD, TRD and WGD, respectively). We found 48 duplicate pairs containing GmPR-1 genes, of which 48%, 21%, 13%, 10% and 8% derived from DD, WGD, TD, TRD, and PD, respectively (Supplementary Table S4). A complementary analysis on the PLAZA server revealed that 95% of the GmPR-1 genes were block duplicates, while 83% were tandem duplicates.

Local duplications (here referred to as tandem and proximal duplications) represent 21% of the GmPR-1 duplicate pairs. However, as genes can be duplicated more than once by different modes, this relative frequency is likely underestimated if each gene is assigned to a single pair and duplication mode. Local duplications explain the physical clustering of 9 GmPR-1 genes on chromosome 13, of *GmPR-1-7* and *GmPR-1-8* within a 6.8 kb region, and of *GmPR-1-9* to *GmPR-1-15* within a 41.6 kb region. Likewise, *GmPR-1-16 to GmPR-1-21* are clustered in a 26.3 kb region on chromosome 15. As a similar trend was observed for AtPR-1 genes (Fig. 2), these findings suggest that independent local duplications have shaped the PR-1 gene repertoire in plants. This physical clustering of defense- and stress-related genes has been described in other species, and it is thought to be an important evolutionary mechanism to ensure coordinated transcriptional regulation [38,45].

The soybean genome has undergone two WGD events, nearly 58 and 13 million years ago, resulting in 75% of its genes being present in multiple copies [46]. As mentioned above, 21% of the GmPR-1 genes were generated by WGD events. Most of these genes diverged near the first WGD (~,58mya) or even in more remote WGDs (Supplementary Table S4). We hypothesize that the ~58mya WGD was important to provide the material for the more recent local duplications that resulted in the current soybean GmPR-1 gene repertoire. In addition, our results suggest that duplicates generated upon the ~13mya WGD were subsequently lost, as there are more WGD pairs from older WGDs than from the ~13mya WGD. We also found that Ka/Ks values were lower than 1 for all pairs derived from all modes of duplication, with median values ranging from 0.13 (TD) to 0.24 (TRD) (Fig. 3d), implying that they are under negative selection. Tandem-derived pairs have the lowest Ka/Ks values and the narrowest Ka/Ks distribution, which indicate that such events happened more recently and around the same time (Fig. 3d).

### 3.5 Biotic and abiotic stresses shaped the cis-regulatory elements of GmPR-1 genes

We identified 13 TF families encoding potential regulators of GmPR-1 genes. The TF families WRKY, NAC, GATA, HD-Zip, ERF and C2H2 had detectable binding domains in the regulatory regions of 11, 4, 3, 3, 2 and 1 GmPR-1 genes, respectively (Fig. 3e). C2H2, HD-Zip and NAC TFs are widely known as key regulators of multiple abiotic stresses in plants, including drought, salinity, cold, and heat stress [47–52]. Likewise, ERFs are ethylene-responsive TFs that have been described as hubs in abiotic stress-related gene regulatory networks [53–55]. GATA TFs have been well described in animals and fungi, but their functions in plants remain unclear. Although plant GATA TFs have been linked to photosynthesis-related pathways [56,57], some studies reported abiotic stress-responsive GATA genes [58,59]. Finally, WRKY TFs are involved in signaling networks activated in abiotic and biotic stresses, particularly in pathogen-associated molecular pattern (PAMP)-triggered immunity [8,60,61]. As the most abundant TF binding sites in GmPR-1 promoters are associated with stress-related TF families, our findings suggest that biotic and abiotic stresses are the major stimuli modulating the transcription of GmPR-1 genes.

### 3.6 Global expression profiles reveal stress-specific and broadly expressed GmPR-1 genes

The global expression profiles of GmPR-1 genes in 1972 RNA-seq samples, comprising multiple tissues and conditions, revealed that 23 of 24 GmPR-1 genes are induced by stress conditions, and *GmPR-1-13* is exclusively expressed under stress (Fig. 4a). Read counts for *GmPR-1-5* are zero or close to zero in almost all samples and greater than 1 in only 17 samples, of which 41% are related to stress conditions. The distribution of Tau indices for tissue specificity indicates that genes from both phylogenetic groups tend to have tissue-specific expression profiles (i.e., Tau values close to 1) (Supplementary Fig. S1).

**Fig. 4.**
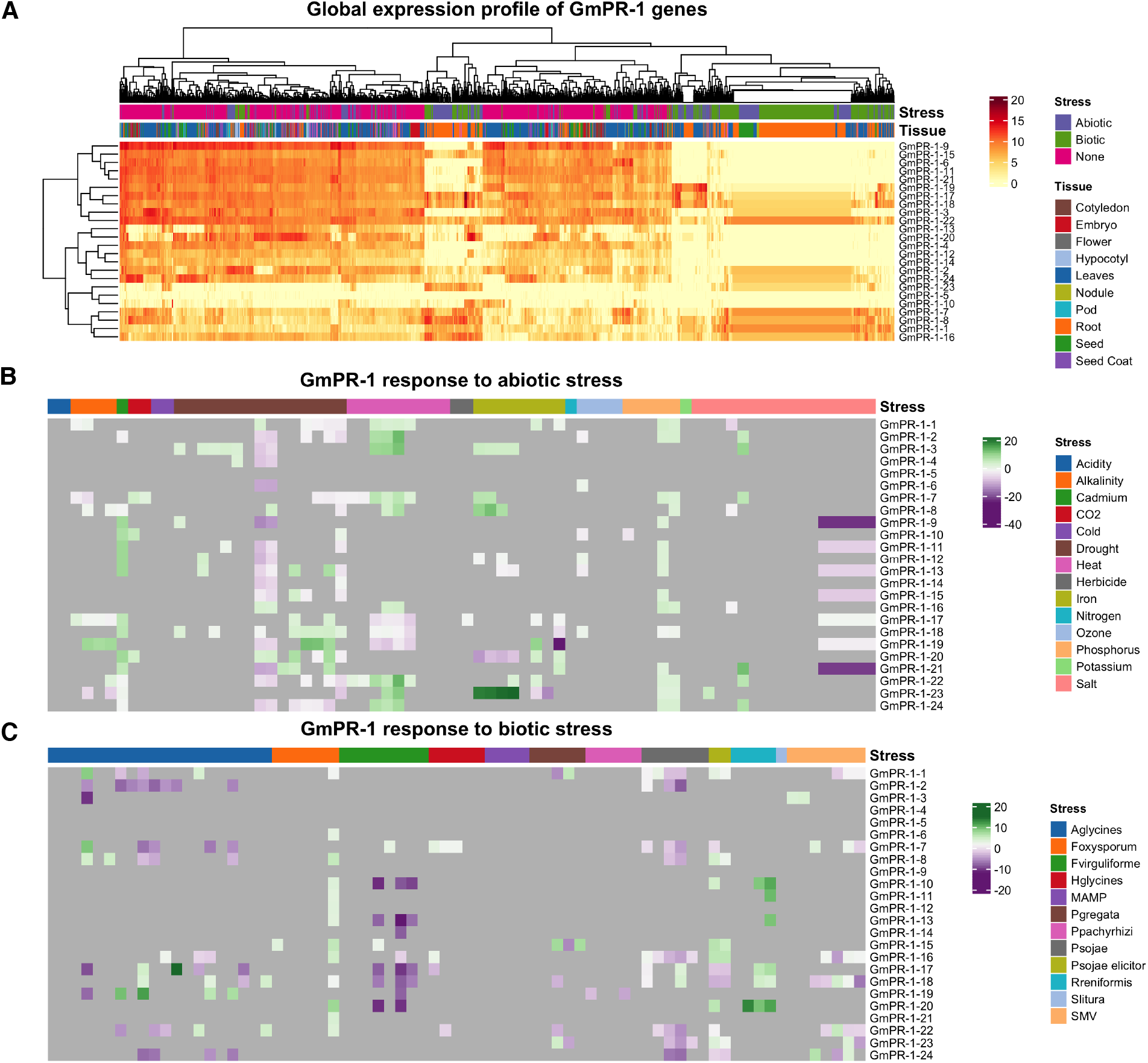
Global and stress-induced expression profiles of GmPR-1 genes. (A) Log_2_ read counts across different tissues and conditions. Dendrograms represent hierarchical clustering of genes and samples. (B) and (C) Log_2_ fold change of GmPR-1 genes in response to abiotic and biotic stress, respectively. Grey represents unchanged expression, green represents up-regulation and purple represents down-regulation. Genes are considered differentially expressed if absolute fold change >1.5 and adjusted *P*-value (Benjamini-Hochberg) <0.05.

PR proteins were originally defined as proteins that are induced only in pathological or related situations [41]. According to van Loon [41], proteins that are homologous to PR proteins but occurred in healthy conditions should be designated as PR-like proteins. Although this classification has been widely overlooked by the scientific community over the years, distinguishing between stress-specific and broadly occurring PR proteins could be useful to identify PR proteins that are exclusively involved in stress response and the ones that can also have endogenous functions. In this sense, *GmPR-1-23* is likely the only gene encoding an exclusively stress-related protein, while the majority of GmPR-1 genes might play a role not only in stress response, but also in plant growth and development. Nevertheless, we hypothesize that a basal level of PR-1 proteins in the absence of stress is required as an immediate first line of response to stress, which would be followed by a stronger, second layer of PR-1 response, harnessed by up-regulation of specific genes.

We found that 86% (21/24) of the GmPR-1 genes are up-regulated under at least one abiotic stress condition, such as alkalinity, salt, cadmium, drought, heat, herbicide, high ozone levels, and iron and phosphorus deficiency (Fig. 4b). Similarly, 20/24 GmPR-1 genes are up-regulated under at least one biotic stress caused by fungi (*Fusarium oxysporum, Fusarium virguliforme, Phialophora gregata,* and *Phakopsora pachyrhizi*), insect (*Aphis glycines*), nematodes (*Heterodera glycines, Rotylenchulus reniformis*), oomycete (*Phytophthora sojae*) and virus (soybean mosaic virus) (Fig. 4c). Interestingly, *GmPR-1-5* and *GmPR-1-14* (CBM- and CAPE-less protein) were not up-regulated under any of the abiotic and biotic stresses.

Further, the oomycete *Phytophthora sojae* and the fungus *Fusarium virguliforme* systematically repressed the expression of GmPR-1 genes, suggesting that these pathogens can down-regulate GmPR-1 transcription. Conversely, *GmPR-1-1*, *GmPR-1-16* and *GmPR-1-18* were up-regulated in the greatest number of biotic stress conditions (*n*=6, 6, 5, respectively), while *GmPR-1-7, GmPR-1-21, GmPR-1-22* were up-regulated in the greatest number of abiotic stress conditions (*n*=5 for all). Thus, these six genes are the most promising genes for biotechnological applications aimed at increasing soybean resistance to biotic and abiotic stresses, respectively.

Little is known about the transcriptional landscape of PR-1 genes in response to various stress conditions, in particular because the studies that investigated the expression profiles of such genes are often limited to a few stress conditions. For instance, PR-1 genes were up-regulated under drought stress in tomato [37] and upon challenge with the fungal pathogen *Septoria nodorum* in wheat [39]. Our findings represent the most comprehensive analysis of the transcriptional responses of PR-1 genes to biotic and abiotic stresses, with 26 different stress conditions. Our results highlight stress-responsive GmPR-1 genes that are potential targets for plant transformation and development of resistant cultivars.

## 4 Conclusions

Here, we reported the soybean PR-1 gene repertoire and characterized it at the sequence, structural and transcriptional levels. The 24 GmPR-1 genes clustered in two phylogenetic groups and are constrained by strong purifying selection. Overall, GmPR-1 proteins are highly conserved in primary and tertiary structures. Further, upstream of GmPR-1 genes, we found multiple binding sites of major stress-related TF families, namely WRKY, HD-ZIP, NAC, ERF, C2H2 and GATA. Finally, we found that the vast majority of GmPR-1 genes are up-regulated under stress conditions, helping uncover candidate genes to be used in the development of soybean lines resistant to both biotic and abiotic stresses.

## Data statement

All data and code used in this study are available in our Figshare repository (DOI: 10.6084/m9.figshare.14330354) to ensure full reproducibility.

## Acknowledgements

This work was supported by Fundação Carlos Chagas Filho de Amparo à Pesquisa do Estado do Rio de Janeiro (FAPERJ; grants E-26/203.309/2016 and E-26/203.014/2018), Coordenação de Aperfeiçoamento de Pessoal de Nível Superior - Brasil (CAPES; Finance Code 001), and Conselho Nacional de Desenvolvimento Científico e Tecnológico. The funding agencies had no role in the design of the study and collection, analysis, and interpretation of data and in writing.

## Author contributions

Conceived the study: FA-S and TMV. Data analysis: FA-S. Funding, project coordination and infrastructure: TMV. Manuscript writing: FA-S and TMV.

