## Supplementary Figures for "Pathogenesis-related protein 1 (PR-1) genes in soybean: genome-wide identification, structural analysis and expression profiling under multiple biotic and abiotic stresses"

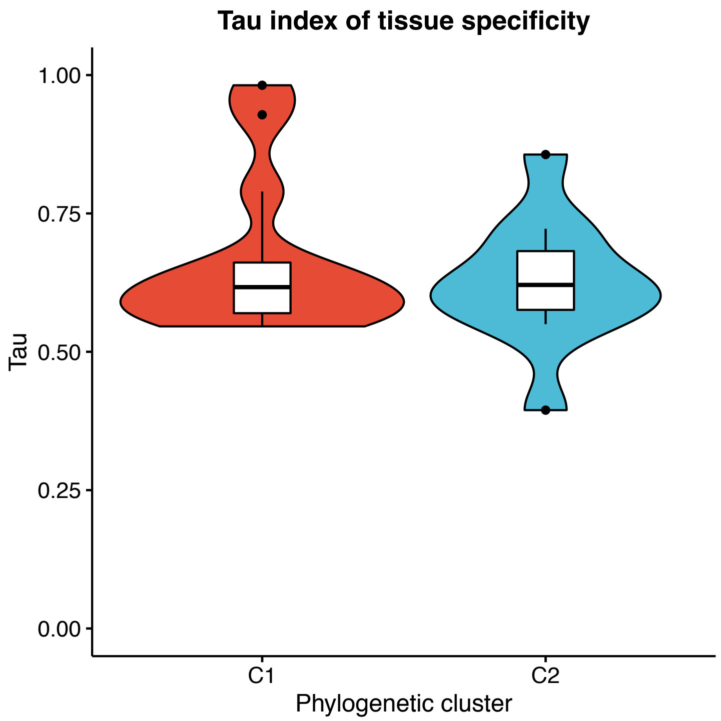


**Fig. 1.** Distribution of Tau indices for phylogenetic clusters 1 and 2. The Tau index is a measure of tissue specificity. Genes with Tau values close to 0 are interpreted as broadly expressed across samples, while Tau values close to 1 indicate tissue-specific expression profiles.
